# Assessing the effects of agricultural intensification on natural habitats and biodiversity in Southern Amazonia

**DOI:** 10.1101/846709

**Authors:** Jan Göpel, Jan Schüngel, Rüdiger Schaldach, Benjamin Stuch, Norman Löbelt

## Abstract

The ongoing trend toward agricultural intensification in Latin America makes it essential to explore intensification measures in combination with assumptions regarding future socio-economic development and policies to protect biodiversity and natural habitats. Information on the future development of land-use and land-cover change (LULCC) under the combination of various driving factors operating at different spatial scale-levels, e.g., local land-use policy and global demands for agricultural commodities is required. The spatially explicit land-use change model LandSHIFT was applied to calculate a set of high-resolution land-use change scenarios for Southern Amazonia. The time frame of the analysis is 2010 - 2030. The resulting maps were analyzed in combination with spatially explicit maps depicting vertebrate species diversity in order to examine the effect of a loss of natural habitats on species ranges as well as the overall LULCC-induced effect on vertebrate diversity as expressed by the Biodiversity Intactness Index in this region. The results of this study indicate a general decrease in Biodiversity Intactness in all investigated scenarios. However, agricultural intensification combined with diversified environmental protection policies show least impact of LULCC on vertebrate species richness and conservation of natural habitats compared to scenarios with low agricultural intensification or scenarios with less effective conservation policies.

## 1. Introduction

Human induced changes to the biosphere have caused severe losses of biodiversity (1, 2). The process of human alteration of natural landscapes and resultant loss of biodiversity is a phenomenon that is mainly attributable to agricultural expansion and intensification. As (3) argue, the growth of agriculture in Brazil has been accompanied by massive deforestation, which is particularly true for the time period from 1970 until the end of the first decade of this century. An area of 18.8% of the original Brazilian Amazon has been deforested since 1970 (4). Land-use dynamics in this time period in the Brazilian Amazon, being distinguished for its biodiversity-rich landscapes (5), have been a major threat to local terrestrial biodiversity due to the conversion of natural ecosystems into cultivated areas (1, 6). Agriculture plays an important role in regard to Brazils GDP (6.1%) (7) and, more importantly, in terms of Brazils exports. A share of 39% of Brazils exported goods are agricultural commodities and products (8). This strong contribution of the agricultural sector to Brazil’s overall economic performance has had positive impacts on social prosperity in the country. According to the (7), the income of 29 million people has been considerably increased, lifting them out of poverty. The inequality (measured by the Gini coefficient) has been lowered by 11% to 0.515. The income level of the poorest 40% rose by 7.1% on average compared to 4.4% income growth of the rest of the population (7). Despite these positive numbers, the global demand for agricultural products is projected to continuously rise over the coming decades (9) driven by global population growth and increasing per capita demand for food, fodder, energy crops, and timber (10, 11). Moreover, changes in food consumption patterns likely further enhance food demands per capita (7). These developments will most likely lead to further expansion and intensification of agricultural area in tropical ecosystems at the expense of natural vegetation and biodiversity (12).

On the one hand, (13) amongst others argue that agricultural intensification (under certain conditions) may be the key for a further increase in productivity, whereby the future destruction of native vegetation can be avoided as far as possible by slowing down the spatial expansion of agriculture e.g. (14–16). On the other hand, some studies argue that agricultural intensification might also lead to area expansion due to the so called “rebound effect” (17) or increasing competiveness of agriculture and, thus higher atainable revenues (18). The latter may only be applicable to situations where commodities with high demand elasticity are involved (19).

Simulation models and scenarios are effective tools to explore current and future land-use changes and to enhance the scientific understanding of land-use change dynamics and their determinants. Many studies exist that examine land-use changes in the Amazon region by employing different models e.g. (20–27). Up to now these models have not been very successful in reproducing the observed land-use changes in the Brazilian Amazon since the end of the 20th century (28). Moreover, several studies assess the impacts of land-use changes on biodiversity in the tropics e.g. (6, 12, 29–32). However, these studies focus on biodiversity in general (6, 12, 32), on a single specific species as an indicator for biodiversity (30, 31, 33) or on a global perspective (29). Thereby, they cannot explicitly assess threats to the regions’ overall vertebrate diversity.

This study aims to address two research questions:

- What will be the effect of a conversion of natural habitats in Mato Grosso (MT) and Pará (PA) on the distribution ranges of vertebrate species?
- What will be the effect of LULCC in MT and PA on vertebrate species diversity measured by the indicator „Biodiversity Intactness Index“?

To address research question 1 we explore the effect of a conversion of natural habitats for the timespan from 2010 to 2030 in a spatial resolution of 900m x 900m. This was accomplished for four socio-economic scenarios and three different taxa (mammals, birds, and amphibians) subdivided into three categories (threatened species, small-ranged species, and endemic species) per taxon. We decided against the inclusion of the category total species richness in our assessment. Total species richness as an indicator for biodiversity can be misleading as it is mainly driven by wide-ranged species (5, 34) which might even be benefit from degraded habitats (35) while especially endemic and small-ranged species are dependent on the intactness of their respective ecosystems (36).

To address research question 2 a pure location-based investigation of future LULCC and its possible impact on vertebrate species diversity is not sufficient. For a more exact analysis an indicator is needed. Therefore, we assessed the change of terrestrial vertebrate species abundance between the reference year 2010 and 2030 as an indicator for the effect of the different scenario assumptions (intensification, extensification, compliance with environmental law, changing consumption pattern) on vertebrate species diversity. The Biodiversity Intactness Index (BII) (37) was calculated for the categories endemic species, small-ranged species, and threatened species for each considered taxon. This indicator provides information to what extent vertebrate species abundance associated with each single grid-cell (900m x 900m) is influenced by LULCC.

## 2. Material and Methods

### 2.1. Study Area

This study focusses on the two Brazilian federal states MT and PA. These states differ greatly in respect to their recent agricultural developments and their level of exploitation of natural habitats due to the Brazilian agricultural development frontier running through this region (38, 39).

PA has an area of 1.25 million km^2^ and a population of 8 million people (40). Only 11,969 km^2^ of the land is used for soybean cultivation (40). In 2015 1,881 km^2^ were deforested which is about the same amount of deforestation as in 2014 (4). The dominant land use sector is cattle ranching with a total herd size of 19.2 million animals (40). A hot spot of LULCC is along the Cuiabá-Santarem highway (BR-163), the most recent of the “development highways” which are used to acquire the agriculturally rather underdeveloped northern parts of Brazil for crop cultivation and cattle ranching (41). The natural vegetation is dominated by dense rainforest (42) covering about 77.6% of the state’s area according to MODIS land cover data (43). More than 40,000 vascular plant species can be found here, of which 30,000 are endemic (42). Over 1,000 bird species are harbored in the Amazon biome (42) as well as a high concentration of mammals, of which many are endemic, especially along the courses of the rivers crossing this biome (5). Of the 875 amphibian species in the country, approximately 50% are concentrated in the Amazon biome (5). Especially here the potential for a loss of vertebrate diversity is high due to a high density of endemic, threatened, and small ranged species (5) as well as ongoing and expected future agricultural expansion (12).

MT has an area of 907,000 km^2^ and a population of 3.2 million inhabitants (40). 69,807 km^2^ of land is used for soybean cultivation (40) and 1,508 km^2^ were deforested in 2015 which constitutes an increase of 16% in comparison to 2014 (4). Another dominant land use sector is cattle ranching with a total herd size of 28.4 million animals (40). Here the expansion of area used for soybean cultivation and cattle ranching could be identified as the primary cause of conversion of natural ecosystems to agricultural land (44, 45). In comparison to PA, MT is more consolidated in terms of agricultural expansion. In recent years, the steep decline of availability of highly productive farmland, policies to curb deforestation and rising land prices have led to a development toward agricultural intensification and away from agricultural expansion (15, 46, 47). MT is covered by two Brazilian biomes, the Brazilian Cerrado and the Amazon rainforest (5, 38). Here, 7,000 plant species, of which 44% are endemic to the Cerrado, can be found. The Cerrado biome is especially rich in bird diversity with 837 species which resembles 49% of all bird species found in Brazil. But also 150 reptile species (50% of all Brazilian reptile species) and 180 amphibian species (28 % endemic to the Cerrado) are found here. The Cerrado biome and its waterways are home to 1,400 fish species, 40% of all fish species occurring in Brazil (48).

### 2.2. Land-use scenarios

In order to explore agricultural intensification and expansion in respect to different socio-economic and policy assumptions, 4 future scenarios have been employed for modeling land use change. These scenarios have been developed during an interdisciplinary research project (CarBioCial; www.carbiocial.de) thematically covering the study area (MT, PA). These scenarios describe plausible future development pathways of Southern Amazonia until the year 2030. Each scenario consists of a qualitative part (storyline) that provides a short narrative of the future world and a set of quantitative information that describe the respective main drivers of LULCC (49, 50). The storylines are elaborated by (51).

The following paragraphs shortly describe the central assumptions of the scenarios. For a comprehensive overview of the quantitative scenario assumptions (crop production, crop yield, population, and livestock) see (52).

The *Trend scenario* assumes a growing demand for agrarian products which is based on an extrapolation of growth trends from 1973 to 2000 specific for each modelled crop. Furthermore, environmental policies like the Brazilian Forest Code or the Soy- and Cattle Moratorium are not part of the assumptions of this scenario. The illegal conversion of natural habitats (protected areas) is prohibited due to good law enforcement. The technological development of agricultural practices in the study area includes an intensification of agricultural production through increasing crop yields. The possibility to intensify pasture management is not considered in this scenario.

Two intensification scenarios (Legal Intensification and Illegal Intensification) assume a growing demand for agrarian products (see *Trend Scenario*) further reinforced by population and GDP growth generated in Asian countries. The technological developments of agricultural practices in the study area include a high degree of agricultural intensification including the intensification of pasture management. The intensification scenarios vary in terms of environmental law enforcement. While the *Legal Intensification scenario* assumes compliance with environmental policies (environmental protected areas, *Brazilian Forest Code*), the *Illegal Intensification scenario* presumes noncompliance with environmental law expressed as the defiance of environmental protected areas concerning agricultural expansion and the noncompliance with the *Brazilian Forest Code*. This scenario assumes the possibility to convert land that is under conservation (e.g. nature reserves), thus opening up spaces that are not allowed for conversion in all other scenarios.

The *Sustainable Development scenario* describes a new social model. This new model includes citizenship, an inclusive economic system, clear land tenure rights, and strong law enforcement including participatory monitoring of deforestation. Furthermore, it portrays a substantial change in terms of anthropogenic consumption pattern, away from a meat oriented diet toward a healthy and sustainable diet as proposed by the WHO (53, 54) including further intensification of crop production. Moreover, the conversion of areas classified as covered by rainforest into agricultural area is not allowed according to the assumptions of the scenario.

### 2.3. Maps of vertebrate diversity and Biodiversity Intactness Index (BII)

We use maps of vertebrate diversity covering the whole area of Brazil (5) to illustrate the overlapping of areas of vertebrate diversity and simulated LULCC in each investigated scenario. The species diversity maps were generated by deriving polygon range data concerning birds from BirdLife International and NatureServe (55) and polygon range data concerning mammals and amphibians from the International Union for the Conservation of Nature (56). These polygon range datasets were rendered at a spatial resolution of 10×10 km in order to produce species diversity maps considering these three groups of terrestrial vertebrates in Brazil (5). These groups were further subdivided into the categories small-ranged species, threatened species, and endemic species. Small-ranged species were defined as those species that have a range smaller than the median for that taxon (2,250,813 km^2^ for birds, 1,230,901 km^2^ for mammals, 66,979 km^2^ for amphibians) in Brazil. For example, a bird species is considered to be small-ranged by occurring naturally in a range of less than 2,250,813 km^2^, which resembles the median distribution range for that taxon in Brazil. Threatened species were defined as vulnerable, endangered, or critically endangered according to the IUCN Red List (57). Endemic species were defined as having at least 90% of their range within Brazil and no part of their range extending more than 50 km beyond the Brazilian border. Overall, 1703 bird species, 637 mammal species and 875 amphibian species were considered in this study.

We calculated the BII accroding to (37)in order to assess the impact of LULCC on overall vertebarte diversity in the time from 2010 to 2030. The BII is defined as the population of a species group *i* under land-use activity *k* in ecosystem *j*, relative to a reference population on the same ecosystem type according to equation 1.

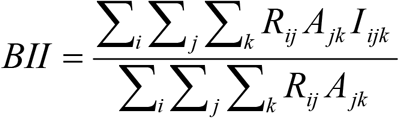

**Equation 1:** Biodiversity Intactness Index (37)

I_ijk_, the *“population impact”*, is the population of a certain species group *i* under land-use activity *k* in ecosystem *j*. A_jk_ is the area of land-use *k* in ecosystem *j*, R_ij_ the number of species of taxon *i* in ecosystem *j*.

Since the calculation is done on grid-cell level, each cell represents an ecosystem. The number of species is the sum of bird species, mammal species, and amphibian species assigned to one cell respectively. In order to formulate the population impact, a combination of impact values from (58, 59) and (60) was employed. These values indicate the reduction of mean species diversity in respect to a certain type of land-cover. The values employed are shown in Table 1. A BII value of 1 indicates a species abundance on the pre-colonial level. An index of 0,5 indicates that the species abundance is reduced by half in reference to the pre-colonial level.

**Table 1:** values used as population impact to calculate BII.

| land-use/ land cover | population | source |
| --- | --- | --- |
| Forest, Barren land | 1.0 | (59) |
| Grassland, Savannah, | 0.94 | (60, 61) |
| Fallow land | 0.5 | (59) |
| Cropland | 0.15 | (59) weighted by proportion of high input agriculture to low input agriculture in Latin America |
| Pasture | Pará: 0.6 /<br>Mato Grosso:<br>0.3 | (58) |
| Mosaic Agricultural Area/rainforest (Legal Reserve) | 0.83* | 20% cropland impact as calculated above and 80% undisturbed forest impact (59) |
| Urban | 0.05 | (59) |

\* A population impact value of 0.83 has been assumed for areas in the Amazon biome that are made up of 20% cropland and 80% rainforest, the so called “Legal Reserve”, in which any kind of deforestation is prohibited according to the *Brazilian Forest Code* (62, 63). This population impact value is considered as an expression for the fragmentation of rainforest.

**Table 2:** aggregation of LandSHIFT land-use classes.

| LandSHIFT land-use types | aggregated land-use types |
| --- | --- |
| evergreen needle forest, evergreen broad-leaved forest, deciduous needle forest, deciduous broad-leaved forest, mixed forest | rainforest |
| closed shrub land, open shrub land | shrub land |
| woody savannah, savannah | savannah (Cerrado) |
| tea, cocoa, coffee, maize, annual oil crops, pulses, rice, tropical roots and tubers, soybean, sugarcane, cassava, wheat | cropland |
| rangeland, pasture | pasture |

A decreasing BII value is an expression for further reduction of biodiversity intactness due to LULCC affecting regions characterized by the occurrence of different species of different taxa. An increasing BII value expresses a recovery of biodiversity intactness mainly due to the displacement of anthropogenic land-use out of these regions or by replacement of certain land-use types by “less harmful” land-use types (e.g. cropland to fallow land) within these regions.

### 2.4. Modeling and assessment protocol

LULCC was analyzed with the spatially explicit LandSHIFT model. The model is fully described in (64) and has been tested in different case studies for Brazil (23, 52, 65). It is based on the concept of land-use systems (66) and couples components that represent the respective anthropogenic and environmental sub-systems. In our study, LULCC is simulated on a raster with the spatial resolution of 900m x 900m that covers the territories of the federal states of MT and PA. The LandSHIFT model generates digital maps for 2010 until 2030 in 5-year time steps that depict the resulting LULCC. For further analysis we aggregated 12 crop types (64) into the land-use class cropland, the 5 distributed forest types (43) into the class rainforest, and the 2 savannah types (43) into the class savannah (Cerrado) according to Table Changes in location and area of the respective land-use types were determined by comparing the maps for 2010 and 2030 using GIS software.

As a second step, we merged the simulated land-use maps of each calculated scenario with maps of vertebrate species diversity regarding three taxa and three categories by overlaying the land-use maps with maps of vertebrate species diversity using GIS software. This promotes quantifying the impact of simulated LULCC on natural habitat area and vertebrate species diversity.

Finally, we calculated the Biodiversity Intactness Index for the reference year 2010 and 2030 according to equation 1. This was accomplished by assigning each land-use type a specific population impact (see Table 1) and multiplying this population impact by the area of the land-use type and the vertebrate species abundance (per taxon and category) associated with that area.

## 3. Results

The main model output comprises time-series of grid maps showing land-use type as well as population and livestock densities. Furthermore, aggregated information on the state-level is produced, including area quantities of each land-use type. Figure 1 shows the land-use and land cover change for each of the scenarios between 2010 and 2030. An elaborate description and discussion of the scenario results can be found in (52).

Figures 2 and 3 present the effect of LULCC driven loss of habitat availability per taxon and category by 2030.

**Figure 1a:**
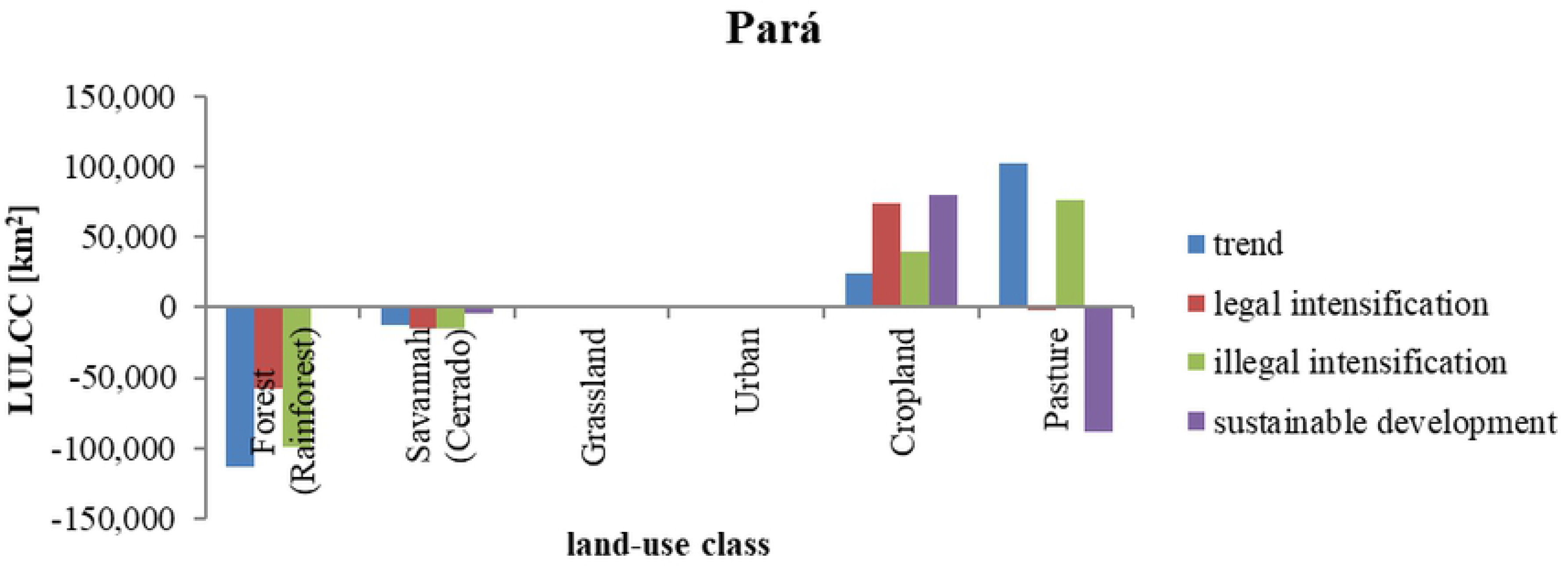
Land-use and land-cover change in Pará between 2010 and 2030.

**Figure 1b:**
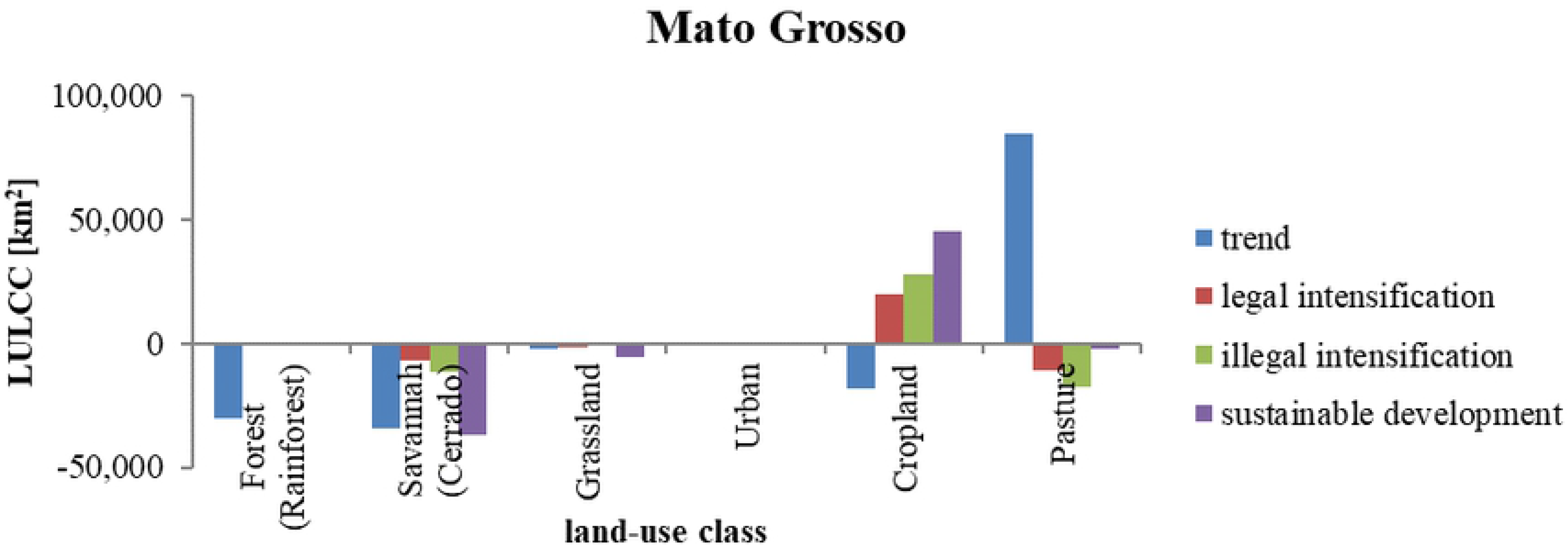
Land-use and land-cover change in Mato Grosso between 2010 and 2030.

**Figure 2:**
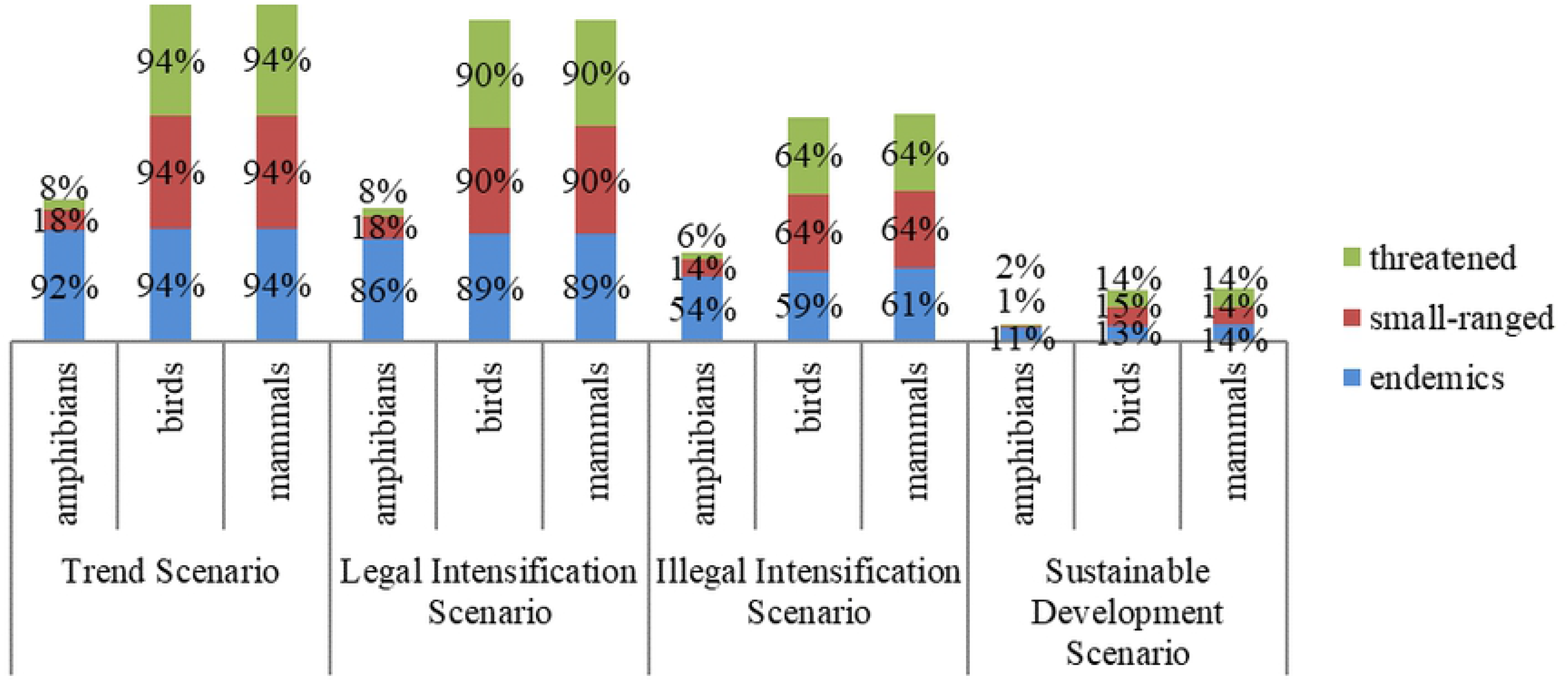
Loss of natural habitat area per assessed taxon and category between 2010 and 2030 in Pará.

**Figure 3:**
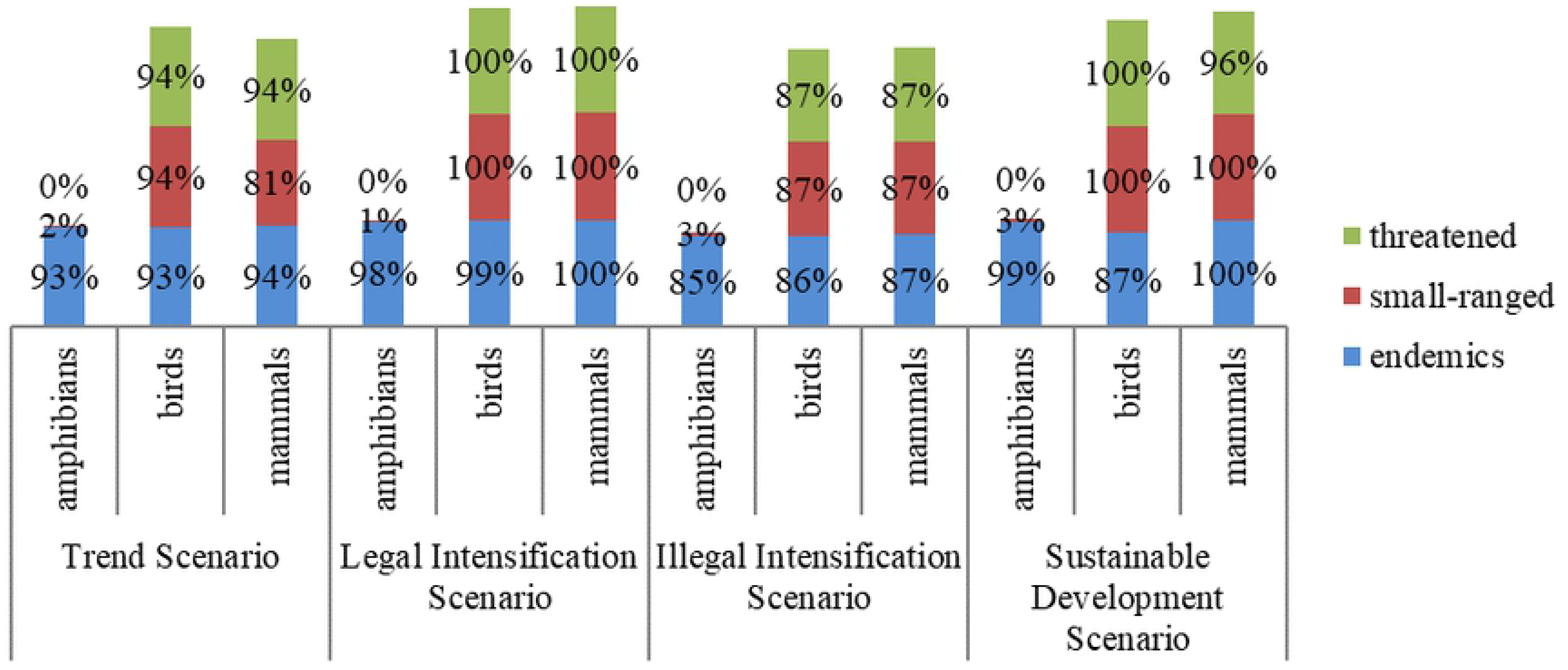
Loss of natural habitat area per assessed taxon and category between 2010 and 2030 in Mato Grosso.

In addition, Tables 3 and 4 present the respective changes in the quantified BII between 2010 and 2030.

**Table 3:** changes of BII in Pará between 2010 and 2030 (Trend=Trend Scenario, LI=Legal Intensification Scenario, ILI=Illegal Intensification Scenario, SD=Sustainable Development Scenario)

|  | taxon | Trend<br>2010 | Trend<br>2030 | rel.<br>Change<br>[%] | LI<br>2030 | rel.<br>Change<br>[%] | ILI<br>2030 | rel.<br>Change<br>[%] | SD<br>2030 | rel.<br>Change<br>[%] |
| --- | --- | --- | --- | --- | --- | --- | --- | --- | --- | --- |
| Amphibians | endemic | 0.79 | 0.71 | -10.1 | 0.71 | -10.1 | 0.73 | -7.6 | 0.79 | 0 |
|  | small-ranged | 0.65 | 0.52 | -20 | 0.53 | -18.5 | 0.57 | -12.3 | 0.66 | 1.5 |
|  | threatened | 0.59 | 0.49 | -16.9 | 0.52 | -11.9 | 0.53 | -10.2 | 0.58 | -1.7 |
| Birds | endemic | 0.8 | 0.71 | -11.3 | 0.71 | -11.3 | 0.73 | -8.8 | 0.8 | 0 |
|  | small-ranged | 0.85 | 0.77 | -9.4 | 0.78 | -8.2 | 0.78 | -8.2 | 0.86 | 1.2 |
|  | threatened | 0.79 | 0.72 | -8.9 | 0.73 | -7.6 | 0.75 | -5.1 | 0.79 | 0 |
| Mammals | endemic | 0.79 | 0.68 | -13.9 | 0.69 | -12.7 | 0.71 | -10.1 | 0.79 | 0 |
|  | small-ranged | 0.86 | 0.77 | -10.5 | 0.78 | -9.3 | 0.79 | -8.1 | 0.86 | 0 |
|  | threatened | 0.79 | 0.65 | -17.7 | 0.69 | -12.7 | 0.72 | -8.9 | 0.78 | -1.3 |

In the following the main LULCC characteristics of each scenario are described. Thereafter, the resultant effect on vertebrate species diversity is addressed by relating natural habitat area loss and vertebrate species diversity as well as by calculating the Biodiversity Intactness Index (Biggs & Scholes, 2005).

### 3.1. Land-use and land-cover change

#### 3.1.1. Pará

In PA, the *Trend Scenario* leads to a loss of tropical rainforests of 113,370 km^2^ (−11.5%), while 12,879 km^2^ (−52.2%) of Cerrado vegetation is converted into urban and agricultural land (Figure 1). The largest fraction of the converted land is used for pasture, which expands by 102,271 km^2^ (+99.6%). Cropland expands by 24,230 km^2^ (+16.4%). In the the *Legal Intensification Scenario*, 57,339 km^2^ (−5.8%) of rainforest and 14,721 km^2^ (−59.7%) of Cerrado vegetation are lost in PA. Cropland increases by 74,717 km^2^ (+50.5%) and pasture areas slightly decrease by 2,238 km^2^ (−2.2%). In regard to the *Illegal Intensification Scenario*, 99,377 km^2^ (−10.1%) of rainforest is converted in PA. Cerrado vegetation decreases by 15,321 km^2^ (−62.2%). Grassland is diminished almost completely. Cropland (+39,181 km^2^, +26.5%) and pasture areas (+76,433 km^2^, +74.4%) increase and cause most loss of natural habitat area. In the *Sustainable Development Scenario,* Cerrado vegetation decreases by 3,711 km^2^ (−15.1%) in PA. No rainforest is converted as rainforest vegetation is fully protected (see Section 2.2). Pasture areas considerably decrease by 89,038 km^2^ (−86.7%). In contrast, cropland increases by 79,676 km^2^ (+53.9%). Most of the newly established cropland area is found in areas that were formerly used for grazing (characterized by relatively high crop yields). Pasture areas decline as a consequence of reduced meat demand in this scenario (see Section 2.2).

#### 3.1.2. Mato Grosso

In the case of the *Trend Scenario* in MT, 34,360 km^2^ (−20.1%) of Cerrado, 30,136 km^2^ (−8.4%) of rainforest and 2,143 km^2^ (−11.1%) of grassland area is affected by LULCC (figure 2). Similar to PA, pasture area expands by 84,588 km^2^ (+50.3%) and is the main driver of the aforementioned changes to natural habitat area. However, contrary to PA, cropland is simulated to decrease by 18,334 km^2^ (−8.4%). The results of the *Legal Intensification Scenario* in MT indicate a loss of 8,937 km^2^ (−4.2%) of Cerrado and 1,750 km^2^ (−9.1%) grassland ecosystems. Rainforest area remains constant. Cropland increases by 19,589 km^2^ (+8.9%), while pasture area decreases by 10,662 km^2^ (−6.3%). In the *Illegal Intensification Scenario*, rainforest cover in MT decreases by only 744 km^2^ (−0.2%); whereas 11,646 km^2^ of Cerrado vegetation (−6.8%) and 1,890 km^2^ (−9.9%) of grassland vegetation is converted. Pasture area decreases by 17,580 km^2^ (−10.5%). Most of the natural vegetation cover is converted to cropland, which expands by 27,939 km^2^ (+12.6%). However, 13,659 km^2^ (77.7%) of the released pasture areas partially accommodate the required expansion of cropland areas. The Sustainable Development Scenario in MT leads to a reduction of Cerrado by 36,731 km^2^ (−21.5%). Grassland vegetation diminishes considerably by 5,761 km^2^ (−30.1%). Rainforest area is protected (see Section 2.2) and remains constant. Cropland expands by 45,092 km^2^ (+20.4%). Pasture area is slightly decreasing by 2,452 km^2^ (−1.5%).

### 3.2. Natural habitat area loss and its effect on vertebrate species diversity

#### 3.2.1. Pará

Figure 2 shows that in PA, the highest impact of a loss of natural habitat area on all taxa was assessed for the *Trend Scenario*. Of the 126,288 km^2^ of converted natural habitats, 94% are characterized by the occurrence of endemic, small-ranged and threatened bird species as well as endemic, small-ranged, and threatened mammal species. However, the picture is a different one when looking at amphibian species. Here 92% of the converted natural habitats are also known for the occurrence of endemic amphibians, but only 18% are known to be a habitat for small-ranged amphibians and only 8% of the converted natural habitats domicile threatened amphibian species. 72,100 km^2^ of natural habitats were converted in the case of the *Legal Intensification Scenario* in PA. Of this area, 90% resemble a habitat for small-ranged and threatened bird and mammal species while 89% are domiciling endemics of the taxa birds and mammals. A share of 86% of the converted area are home to endemic amphibians while 18% are known habitats of small-ranged amphibians and only 8% domicile threatened amphibians. In the case of the *Illegal Intensification Scenario*, 114,939 km^2^ of natural habitat area is lost. Of this area, 59% are home to endemic bird species while 64% domicile small-ranged and threatened species of this taxon. Concerning mammal species, 61% of the converted natural habitat area domicile endemic species and 64% are known habitats of small-ranged and threatened mammal species. Amphibian species are less disturbed as 54% of the lost natural habitats are home to endemic amphibian species while 14% and 6% are domiciling small-ranged and threatened amphibian species respectively. The least negative effect on vertebrate species diversity due to a conversion of natural habitats is discernible in the case of the *Sustainable Development Scenario*. Not only the area of converted natrual habitats is less as compared to the other scenarios, also the share of this area that is a known habitat to vertebrate species is relatively low. Here, 14% of the 3,752 km^2^ are home to endemic, small-ranged and threatened mammals. 13% are domiciling endemic birds, 15% are habitats of small-ranged birds and 14% domicile threatened bird species. 11% house endemic amphibians, while only 1% and 2% domicile small-ranged and threatened amphibians respectively.

#### 3.2.2. Mato Grosso

As Figure 3 shows, the assumptions made for the *Trend Scenario* in MT result in a strong disturbance of vertebrate species diversity. An area of 66,634 km^2^ of natural habitats is converted of which 93% are home to endemic bird species while 94% domicile small-ranged, and threatened bird species. 94% of the converted area is home to endemic and threatened mammals while 81% of the converted natural habitat area is a habitat for small-ranged mammals species. 93% of the converted natural habitat area are home to endemic amphibians while only 2% and 0% of this area domicile small-ranged and threatened amphibian species respectively. The latter may not be consistent with the actual situation as threatened amphibians are especially data deficient in Brazil (5). In the case of the *Legal Intensification Scenario*, 100% of the converted natural habitats (8,910 km^2^) are domiciling small-ranged and threatened birds as well as endemic, small-ranged, and threatened mammals. 99% of the lost natural habitat area is home to endemic birds. 98% are housing endemic amphibians while only 1% are known habitats of small-ranged amphibians. Due to data deficiencies regarding threatened amphibians in Brazil, no threatened amphibians species are spatialized to the converted natural habitat area. Concerning the *Illegal Intensification Scenario*, 86% of the converted natural habitats (14,280 km^2^) are considered distributions range of endemic bird species while 87% domicile small-ranged and threatened bird species respectively. Also 87% of the converted natural habitats domicile endemic, small-ranged, and threatened mammals. 85% of the converted natural area shelter endemic amphibians while only 3% are home to small-ranged amphibians. Again, no data on threatened amphibians was available in regard to the converted area in the case of the *Illegal Intensification Scenario*. The assumptions made in the case of the *Sustainable Development Scenario* lead to a reduction of natural habitats by 42,492 km^2^. 88% of that area are known habitats of endemic bird species while 100% shelter small-ranged and threatened bird species. Also 100% of the converted natural habitat area domicile endemic and small-ranged mammals while 96% of the lost natural area are home to threatened mammals. 99% of the converted natural habitat area shelter endemic amphibians and only 3% are home to small-ranged amphibians. For the mentioned reasons no data concerning threatened amphibians was available for the converted natural habitats in MT.

### 3.3. Biodiversity Intactness Index

#### 3.3.1. Pará

Table 3 shows, the impact on species diversity, as expressed by changes of the BII, is strongest as calculated in the case of the *Trend Scenario* in PA. We found especially strong redcutions of BII for threatened mammals, with a reduction of −17.7%, followed by threatened amphibians with a reduction of −16.9%, and endemic mammal species with a reduction of - 13.9%. In the case of the *Legal Intensification Scenario*, we see BII value decreases for all taxa and categories. Especially strong disturbances can be discerned in the case of small-ranged amphibian species (−18.5%), endemic mammals as well as threatened mammals with - 12.7% respectively. In the case of the Illegal Intensifcation Scenario especially small-ranged mammal species (−12.3%), endemic mammals (−10.1%), and threatened amphibian species (−10.1%) were strongly affected. Interestingly, the negative effect in the case of the *Illegal Intensification Scenario* is lower compared to the *Legal Intensification Scenario*. The negative effects of forest fragmentation has a stronger negative impact on BII than the opening up of former protected areas for agricultural expansion. The lowest negative effect was simulated in the case of the Sustainable Development Scenario. The highest decrease was calculated for threatened amphibians (−1.7%) while the BII value for threatened mammals decreased by 1.3%. All other BII values remained constant or even increased as was the case for small-ranged amphibians (+1.5%) and small-ranged bird species (+1.2%).

#### 3.3.2. Mato Grosso

Table 4 shows, the impact on species diversity in MT is more moderate in the case of the *Trend Scenario* compared to the situation in PA. Although, one has to keep in mind that the BII values in MT are on average 0.14 points below those calculated for PA due to MT being more consolidated in agricultural terms. The highest reduction of BII in Mato Grosso was simulated in the case of the *Trend Scenario*. Here, threatened bird species (−13.6%), threatened mammals (−10.3%), and endemic mammals (−9.5%) are especially affected. In the case of the Legal Intensification Scenrio, we see an decreasing BII value for all taxa and categories with the exception of small-ranged birds which remains constant. We found especially strong decreases for threatened mammals (−7.4%) and threatened birds (−6.8%). Concerning the *Illegal Intensification Scenario*, we see decreasing BII values for all taxa and categories. Here especially small-ranged amphibians (−35.7%), threatened mammals (−11.8%) and threatened bird species (−15.3%) are strongest impacted. This effect can be explained by the opening up of former protected areas for agricultural expansion. Especially the Pantanal, known for its species richness in regard to birds and amphibians (Figuera et al. 2006; Jenkins et al. 2015), is affected by the displacement of agricultural areas into formerly protected areas. Interestingly, the Sustainable Development Scenario in MT results in a strong negative impact. It becomes obvious that all taxa and categories are affected negatively, especially endemic amphibians (−7.5%) and endemic birds (−7.6%) as well as threatened birds (−10.2%) and threatened mammals (−7.4%).

**Table 4:** changes of BII in Mato Grosso between 2010 and 2030 (Trend=Trend Scenario, LI=Legal Intensification Scenario, ILI=Illegal Intensification Scenario, SD=Sustainable Development Scenario)

|  | taxon | Trend<br>2010 | Trend<br>2030 | rel.<br>Change<br>[%] | LI<br>2030 | rel.<br>Change<br>[%] | ILI<br>2030 | rel.<br>Change<br>[%] | SD<br>2030 | rel.<br>Change<br>[%] |
| --- | --- | --- | --- | --- | --- | --- | --- | --- | --- | --- |
| Amphibians | Endemic | 0.67 | 0.62 | -7.6 | 0.66 | -1.5 | 0.67 | 0 | 0.62 | -7.5 |
|  | Small-ranged | 0.56 | 0.54 | -3.6 | 0.55 | -1.8 | 0.36 | -35.7 | 0.54 | -3.6 |
|  | Threatened | n.a. | n.a. | n.a. | n.a. | n.a. | n.a. | n.a. | n.a. | n.a. |
| Birds | Endemic | 0.66 | 0.6 | -9.1 | 0.65 | -1.5 | 0.67 | 1.5 | 0.61 | -7.6 |
|  | Small-ranged | 0.62 | 0.57 | -8.1 | 0.62 | 0 | 0.6 | -3.2 | 0.59 | -4.8 |
|  | Threatened | 0.59 | 0.51 | -13.6 | 0.55 | -6.8 | 0.5 | -15.2 | 0.53 | -10.2 |
| Mammals | Endemic | 0.63 | 0.57 | -9.5 | 0.62 | -1.6 | 0.64 | 1.6 | 0.59 | -6.4 |
|  | Small-ranged | 0.66 | 0.6 | -9.1 | 0.65 | -1.5 | 0.64 | -3.0 | 0.63 | -4.6 |
|  | Threatened | 0.68 | 0.61 | -10.3 | 0.63 | -7.4 | 0.6 | -11.8 | 0.63 | -7.3 |

## 4. Discussion

Agricultural intensification has played an important role in regard to recent agricultural production growth in Brazil and is likely to further increase Braziĺs crop and beef production considerably (67). The observed decoupling of production increases from deforestation in the latter half of the first decade of this century (20, 68) have shown that the intensification of agricultural systems not only supports food provisioning, it also limits the expansion of agricultural area; thus the destruction of natural habitats (69).

This trend is confirmed in our study, which is among the first to investigate the impact of projected LULCC on a proxy for overall terrestrial vertebrate diversity on a regional scale. The negative effect of projected future agricultural production growth on natural habitats and vertebrate species diversity is considerably reduced through agricultural intensification and particularly through intensification of grazing intensities on pastures (compare with (14, 69)). The two agricultural intensification scenarios show substantially less LULCC compared to the *Trend Scenario* based on constant crop yields and grazing intensities of the reference year 2010.

In PA, the loss of natural habitats could be reduced by 43% in the case of the *Legal Intensification Scenario* and by 9% in case of the *Illegal Intensification Scenario* compared to the *Trend Scenario*. This is also confirmed by (15) who have shown that the encouragement of an intensification of pastures, either through subsidies of intensified systems or tax on extensive pastures, considerably limits the conversion of natural habitats until 2020. Also, in these two intensification scenarios the share of the converted natural habitats that is known as habitats to vertebrate species is less as in the case of the *Trend Scenario* (see Figure 2). The strongest effect in regard to mitigating the effect of a conversion of natural habitats on vertebrate species diversity could be achieved in the Sustainable Development Scenario. Here, the loss of natural habitats could be reduced by 97% (from 126,288 km^2^ to 3,752 km^2^) in comparison with the *Trend Scenario*. Additionally, only 11-15% of the converted 3,752 km^2^ natural areas in the Sustainable Development Scenario are known habitats of amphibian, bird, and mammal species.

In MT, the area of affected natural habitats in the case of the *Illegal Intensification scenario* could be reduced by 78% (from 66,634 km^2^ to 14,280 km^2^) compared to the *Trend Scenario* with 85-87% of that area domiciling vertebrate species diversity. This reduction due to agricultural intensification is surpassed by the *Legal Intensification Scenario*. Thereby, the loss of natural habitats could be limited by 86% (from 66,634 km^2^ to 8,914 km^2^) compared to the *Trend Scenario* with the caveat that 100% of that area is habitat to amphibian, bird, and mammal species. These results suggest that intensification measures area especially effective if combined with adequate conservation policies as assumed in the case of the *Legal Intensification Scenario* (see Section 2.2). This is confirmed by several other authors e.g. (70, 71) who found that the intensification of agricultural production and the protection of natural habitats have the highest impact in terms of limiting the conversion of natural habitats, and thus promoting the conservation of vertebrate species diversity. The optimal combination of intensification and conservation measures in terms of a maximum reduction of converted natural habitats depends on the present situation in the respective region as the heterogeneity of losses of natural habitats in PA and MT under the respective scenario assumptions illustrates. Concerning the Sustainable Development Scenario in MT, cropland area expands especially strong due to a shift of anthropogenic consumption patterns away from meat toward crop intake while there is only a slight release of pasture area,. Therefore, the decrease of pasture area can only partially counteract the expansion of cropland, and thus the loss of natural habitat area. Overall, this leads to a reduction of converted natural habitat area by 36% compared to the *Trend Scenario* (see Figure 1). This is considerably less as compared to both intensification scenarios.

The effects of a loss of natural habitats on vertebrate species diversity are confirmed by our assessment of the BII in PA. We see decreasing BII values in the case of all simulated scenarios for all assessed taxa and categories with few exceptions (small-ranged birds and amphibians in regard to the Sustainable Development Scenario) in PA. These decreases are especially strong in the case of the *Trend Scenario* and both intensification scenarios. In the case of the *Legal Intensification Scenario*, the requirement to establish a “Legal Reserve”, a share of 80% of any holding covered by rainforest that needs to be preserved, leads to a fragmentation and consequently degradation of rainforest habitats e.g. (72, 73). This fragmentation in turn causes a considerable reduction of the BII values in the case of the *Legal Intensification Scenario* across all assessed taxa and categories which are stronger as in the *Illegal Intensification Scenario* where a fragmentation of rainforest habitats is not assumed. The negative effect of LULCC on vertebrate species diversity as expressed by the BII is weakest in the case of the Sustainable Development Scenario. This is attributable to the effect of a substantial reduction of the global meat intake which leads to a significant reduction of pasture area, and thus overall expansion of agricultural area into natural habitats. Even without an intensification of pasture management, this reduction of pasture area is able to compensate further expansion of cropland area. Thereby, cropland expansion is limited to released pasture areas, which mitigates LULCC pressure on natural habitats.

The positive implication of agricultural intensification on biodiversity found in PA is confirmed also in MT. Here, the BII values decrease in almost all assessed taxa and categories in the case of the *Legal Intensification Scenario* and *Illegal Intensification Scenario*. This can be explained by the reduced land requirements in both scenarios. As current protected areas have no legal conservation status in the *Illegal Intensification Scenario*, loss of natural habitats is simulated in the biodiverse areas of the Pantanal. Here, (5) found especially strong concentrations of bird and amphibian species. This explains why we measure especially strong decreasing BII values for all threatened vertebrate species as well as small-ranged amphibian species in the *Illegal Intensification Scenario* in MT. In contrast, current protected areas are assumed being effectively conserved in the *Legal Intensification Scenario*, which displaces LULCC from the Pantanal to other not conserved, less biodiverse areas. This prevents strongly decreasing BII value for threatened vertebrate species and especially small-ranged amphibian species in the *Legal Intensification scenario* in MT. Moreover, the overall higher BII values in the *Legal Intensification Scenario*, compared to the *Illegal Intensification Scenario*, shows that effective conservation of existing protected areas can further enhance biodiversity in MT in 2030. (29) calculated BII values of around 70% for the Brazilian Cerrado (mainly located in MT) as well as 85% for the Amazon biome (mainly located in PA). This agrees well with our calculation for the year 2010 of 0.59-0.68 (MT) and 0.65-0.86 (PA) respectively (see Tables 3 and 4.) The fact that the estimates found in our study are slightly lower than those estimated by (29) is explained by taking into account that Newbold et al. focused on the whole Cerrado and Amzonia region while we assess a subregion that is and was characterized by especially strong LULCC dynamics.

On the one hand, the potential for agricultural intensification in the Amazon may hint at the way of sustaining food production here (13) but, on the other hand, it also draws a distressing picture of the future in regard to negative impacts of intensification measures (74). Despite all the positives of an intensification of agricultural production in regard to a conservation of natural habitats, the negative impacts of an intensified agriculture cannot be neglected. Especially pesticide, herbicide, and fertilizer application have to increase in order to increase grass- and cropland productivity (75). The increased application of such products will have negative effects on biodiversity (74). Especially the use of pesticides in tropical regions has strong negative effects on amphibian populations because they are more susceptible to pesticide use as compared to amphibian populations in temperate regions (76). Therefore, biodiversity on intensified cropland is likely to decrease. Furthermore, the adoption of intensified agricultural production will cause higher costs of production. These higher costs may hinder smallholder farmers to apply such techniques. This in turn will imperil their ability to stay competitive in comparison to large land holders who have better access to monetary resources and can make larger investments into the intensification of agricultural production (77). An increased livestock production in intensified systems (especially feedstock systems) will increase the demand for livestock fodder production which, in turn, will induce an expansion of cropland area and may be a cause of additional deforestation (77).

### 4.1. Limitations and uncertainties of the study

Concerning the data that was used to assess the effect of a loss of natural habitat area on species diversity as well as the BII values, the species diversity maps (5), issues of data deficiency have an impact on our estimates. Especially amphibian and mammal species are understudied. Data deficient mammals are mainly concentrated in the Amazon whereas around 30% of all assessed amphibians are generally data deficient (5). This may lead to an underestimation of the impact of loss of natural habitats on vertebrate diversity especially in regions covered by rainforest (Amazon). Notable are threatened amphibian species. Here, only 4% of the assessed species appear to be threatened, whereas the global rate of threatened amphibians lies at 31% (78) suggesting that the high data deficiency in regard to this taxon and the investigated area are significantly influencing our results.

Moreover, we do not holistically explore the effects of agricultural intensification on natural habitats and its biodiversity. In order to do so it would require an analysis of all factors of agricultural intensification that positively or negatively influences wildlife and habitats. This analysis would have to include emissions caused due to intensification (livestock, fertilizers etc.) and their effect on biodiversity as well as indirect LULCC, for instance caused by a cropland expansion due to an increasing demand for fodder in feedstock systems. The inclusion of these factors was beyond the scope of this study.

In all of our scenarios we assume that increases of crop yields until 2030 can be achieved by technological improvements and a more intensive agricultural management alone. At the same time studies such as (65, 79) point out that climate change might have negative effects on crop yields in Amazonia. It is important to note that this situation might occur until the mid or end of this century when changes in temperature and precipitation are projected to become more intense (80) with potentially stronger negative impacts on crop yields e.g. (79, 81).

